# Conformational transitions of the HIV-1 Gag polyprotein upon multimerization and gRNA binding

**DOI:** 10.1101/2023.08.16.553549

**Authors:** Puja Banerjee, Gregory A. Voth

**Affiliations:** Department of Chemistry, Chicago Center for Theoretical Chemistry, Institute for Biophysical Dynamics, and James Franck Institute, The University of Chicago, Chicago, IL 60637

**Keywords:** HIV-1 Gag assembly, multi-domain protein, protein-protein interactions, Gag multimerization, peripheral membrane proteins

## Abstract

During the HIV-1 assembly process, the Gag polyprotein multimerizes at the producer cell plasma membrane, resulting in the formation of spherical immature virus particles. Gag-gRNA interactions play a crucial role in the multimerization process, which is yet to be fully understood. We have performed large-scale all-atom molecular dynamics simulations of membrane-bound full-length Gag dimer, hexamer, and 18-mer. The inter-domain dynamic correlation of Gag, quantified by the heterogeneous elastic network model (hENM) applied to the simulated trajectories, is observed to be altered by implicit gRNA binding, as well as the multimerization state of the Gag. The lateral dynamics of our simulated membrane-bound Gag proteins, with and without gRNA binding, agree with prior experimental data and help to validate our simulation models and methods. The gRNA binding is observed to impact mainly the SP1 domain of the 18-mer and the MA-CA linker domain of the hexamer. In the absence of gRNA binding, the independent dynamical motion of the NC domain results in a collapsed state of the dimeric Gag. Unlike stable SP1 helices in the six-helix bundle, without IP6 binding, the SP1 domain undergoes a spontaneous helix-to-coil transition in the dimeric Gag. Together, our findings reveal conformational switches of Gag at different stages of the multimerization process and predict that the gRNA binding reinforces an efficient binding surface of Gag for multimerization, as well as regulates the dynamic organization of the local membrane region itself.

**Significance:** Gag(Pr_55_^Gag^) polyprotein orchestrates many essential events in HIV-1 assembly, including packaging of the genomic RNA (gRNA) in the immature virion. Although various experimental techniques, such as cryo-ET, X-ray, and NMR, have revealed structural properties of individual domains in the immature Gag clusters, structural and biophysical characterization of a full-length Gag molecule remains a challenge for existing experimental techniques. Using atomistic molecular dynamics simulations of the different model systems of Gag polyprotein, we present here a detailed structural characterization of Gag molecules in different multimerization states and interrogate the synergy between Gag-Gag, Gag-membrane, and Gag-gRNA interactions during the viral assembly process.

## Introduction

Human Immunodeficiency Virus type-1 (HIV-1) virus particle assembly in the infected cells is a complex process involving a highly regulated network of several specific and non-specific protein: protein, protein: lipid, protein: RNA interactions, and the formation of multi-protein complexes (1–3). The major structural component of HIV-1 is the 55 kDa multidomain protein, Gag, that mainly drives the viral assembly process. This 500 amino acid protein comprises several independently folded structural subdomains, connected by flexible linkers (4). Approximately 2500 Gag molecules assemble at the plasma membrane (PM) of the infected cell around a dimer of full-length genomic RNA (gRNA) forming a spherical, non-infectious immature virus particle (2,5–9). The myristoylated matrix (Myr-MA) domain mediates membrane binding of Gag and envelope (Env) glycoprotein incorporation into the virions (10–21). Apart from MA, the other domains are capsid (CA), spacer peptide-1 (SP1), nucleocapsid (NC), spacer peptide-2 (SP2), and p6, in order of appearance in the protein sequence (***Figure 1***). The CA and NC domains play crucial roles in the Gag multimerization process (22) and selective packaging of the gRNA (23), while the p6 domain primarily mediates the viral budding (24).

**Figure 1:**
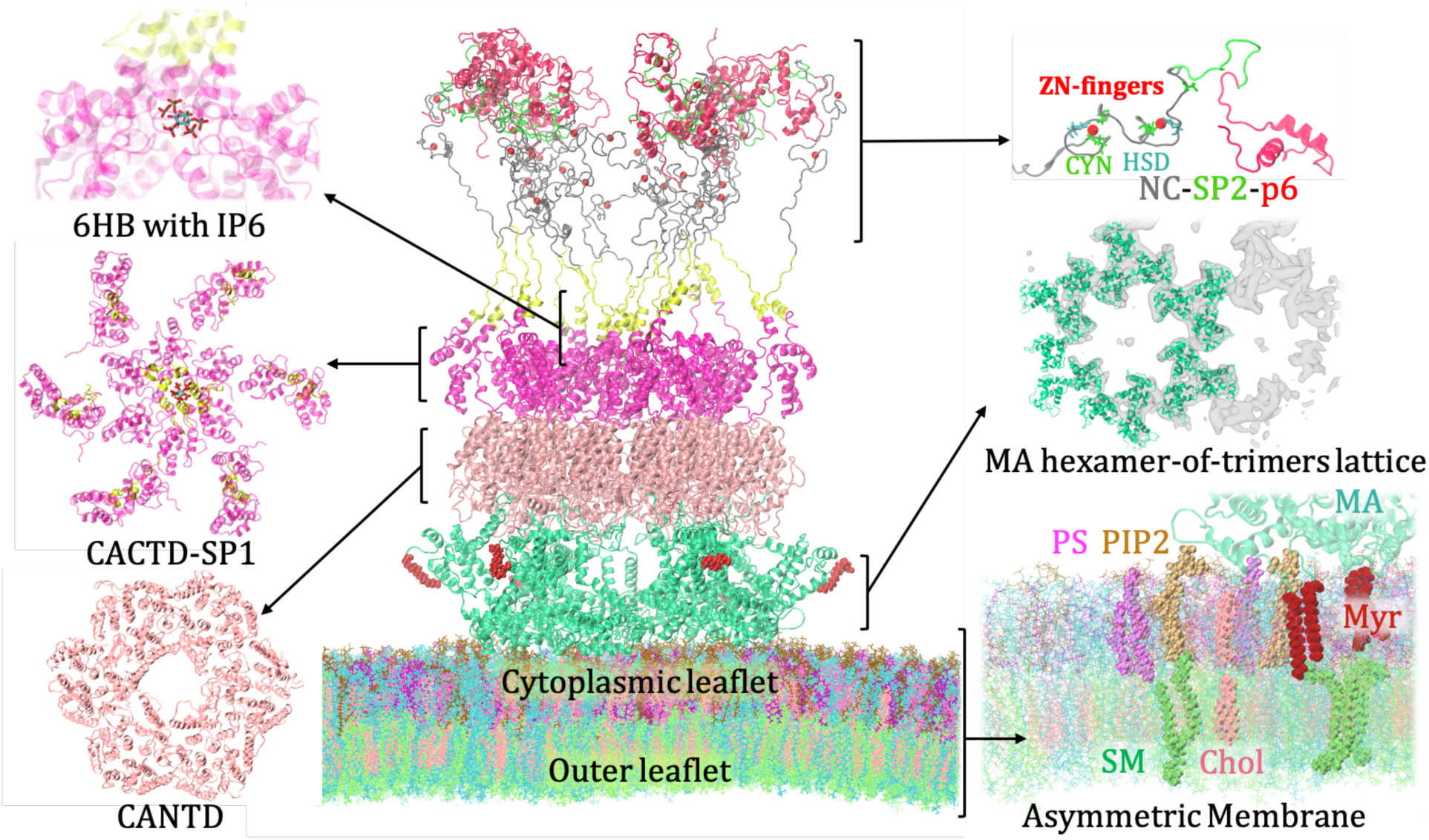
Membrane-bound immature HIV-1 Gag bundle. The final structure of membrane-bound full-length Gag 18-mer from a 1 μs AAMD simulation trajectory. The atomic model of full-length Gag 18-mer was built using experimental structural data of MA hexamer-of-trimers (green), CASP1 oligomers (pink and yellow), NMR structure of NC domain(gray) with CCHC ZN finger motifs, and p6 domain (red). IP6 anion binds to the six-helix bundle (6HB) and confers stability to the inner hexamer. Myristoyl groups (red) attached to the N-terminus of the MA domain anchor the inner leaflet of the bilayer, enriched in anionic phospholipids, such as PIP2 and PS.

Cryo-ET, X-ray crystallography, and NMR spectroscopy have revealed atomic structures of individual domains in their monomeric (NC and p6) and multimeric (MA, CA) states (22,25–28). Although full-length Gag protein has been successfully isolated in soluble dimeric or trimeric form, the atomic structures could not be resolved due to its flexible nature (29–31). Earlier, in a collaborative study with Briggs and coworkers, using cryo-ET data and all-atom molecular dynamics (AAMD) simulations, the Gag dimers were identified as the primary assembly unit that is recruited into the immature growing lattice, mediated by the intra-hexameric interactions of the CA_CTD_ domain (32). Gag-Gag interactions during the assembly process are predominantly triggered by the hexameric interactions mediated by the major homology region (MHR, residues 285-304) of the CA_CTD_ domain and SP1. The last eight residues of the CA_CTD_ domain and the first eight residues of the SP1 domain (residues 356 to 371) form a six-helix bundle (6HB) (22,32–36). X-ray crystallography and cryo-EM maps reveal that the 6HB structure is stabilized by the cellular metabolite inositol hexakisphosphate (IP6) as well as the hydrophobic knobs-in-holes interactions (34,36,37). The polyanion, IP6, binds the rings of basic LYS residues (LYS290, LYS359) and the mutations of either LYS residues have been observed to disrupt immature viral particle production and infectivity (37,38). A recent coarse-grained molecular dynamics (CGMD) study revealed that other than the role of IP6 as an assembly accelerator, it promotes kinetically-trapped fissure-like defects in the immature lattice which might play an important role in the later stages of the viral replication cycle (39). In addition, homo-dimeric and homo-trimeric interactions are mediated by the CA_NTD_ and CA_CTD_ residues. Helix1 of CA_NTD_ and helix 9 of CA_CTD_ domains form dimeric contacts and helix 2 of CA_NTD_ stabilizes trimeric interfaces between adjacent Gag molecules (22,33,34). Furthermore, the CA_CTD_-SP1 domain is an established target for maturation inhibitors, such as Bevirimat (BVM) and PF-46396 (PF-96) (40). BVM stabilizes 6HB and inhibits the last proteolytic cleavage event of Gag between CA and SP1 (34,36). These latter insights have recently been re-emphasized in additional combined experimental-computational studies (41). The dynamics of Gag molecules play a crucial role in the assembly, budding, and maturation process. Several studies interrogated Gag dynamics in the lattice (42) and during the early stage of the viral assembly (43). Using the reaction-diffusion model, Guo et al. predicted the timescale for lattice dynamics and protease dimerization (44).

The membrane-bound Myr-MA domain assembles as hexamer of trimers in the immature virions (25). Membrane binding of the MA domain is predominantly mediated by the lipid-specific interactions with a minor phospholipid component of the plasma membrane, phosphatidylinositol 4,5-bisphosphate (or PIP2) (16–20). Experiments using a combination of super-resolution stimulated emission depletion (STED) microscopy and fluorescence correlation spectroscopy (STED-FCS) have suggested that HIV-1 Gag selectively traps PIP2 and cholesterol to create specialized membrane microdomains for viral assembly, but not phosphatidylethanolamine (PE) and sphingomyelin (SM) (45). Recently, using fluorescence quenching assays, Wen *et al.* demonstrated that Gag multimerization promotes PIP2 clustering(46). An enrichment of PIP2 has further been identified in the viral membrane compared to the host-cell plasma membrane (47). Besides MA-PIP2-specific interactions, non-specific electrostatic interactions with anionic phospholipids such as phosphatidylserine (PS), and hydrophobic interactions between membrane-inserted N-terminal myristoyl (Myr) group and lipid tails of the membrane bilayer stabilize MA-membrane binding (21,48,49). The Myr group remains in equilibrium between a sequestered conformation being buried into the hydrophobic pocket of MA, and an exposed conformation. NMR experiments have suggested that trimerization of MA facilitates the transition between sequestered to exposed state, known as the “myristoyl switch” (15).

Apart from the protein-protein and protein-membrane interactions described above, the HIV-1 assembly process involves the packaging of viral gRNA mediated primarily by the specific interactions between nucleocapsid (NC) domain with the 5’ untranslated region (UTR) of gRNA, which contains the Ψ packaging signal (43,50–54). The NC domain contains two CCHC-type zinc-finger (ZF) motifs, between them N-terminal ZF predominantly drives the gRNA recognition (55–57). An experimental study using a hybrid fluorescent proximity/sedimentation velocity method in combination with calorimetric analyses suggested that gRNA binding promotes Gag dimerization which is distinct and stronger than CA or SP1 domain-mediated oligomerization (58). Another recent study interrogated multiple factors that impact genome packaging using a fluorescence microscopy-based single virion analysis technique and suggested that gRNA binding is intricately related to Gag multimerization and membrane binding (59). In that study, mutants with defective multimerization or plasma membrane anchoring were observed to affect gRNA packaging. Using coarse-grained simulations, a prior study from our group has demonstrated that both gRNA and membrane binding act as scaffolds that promote Gag assembly (60). While it is evident that Gag-Gag, Gag-membrane, and Gag-gRNA interactions play an important role in the viral assembly process, significant knowledge gaps remain in our understanding regarding the impact of Gag-gRNA interactions on the structure and dynamics of the membrane-bound Gag polyproteins.

In the present study, we have carried out large-scale all-atom molecular dynamics (AAMD) simulations of full-length Gag proteins in different multimerization states, bound to an asymmetric lipid membrane. Four-site CG models of Gag monomers are then parametrized using the EDCG (Essential Dynamics Coarse Graining) mapping scheme (61) and the hENM (heterogeneous Elastic Network Model) (62) to assess interdomain interactions of Gag in different systems (**Figure 2**). Our findings also provide new insights about the disordered linker domains which remained largely unknown.

**Figure 2:**
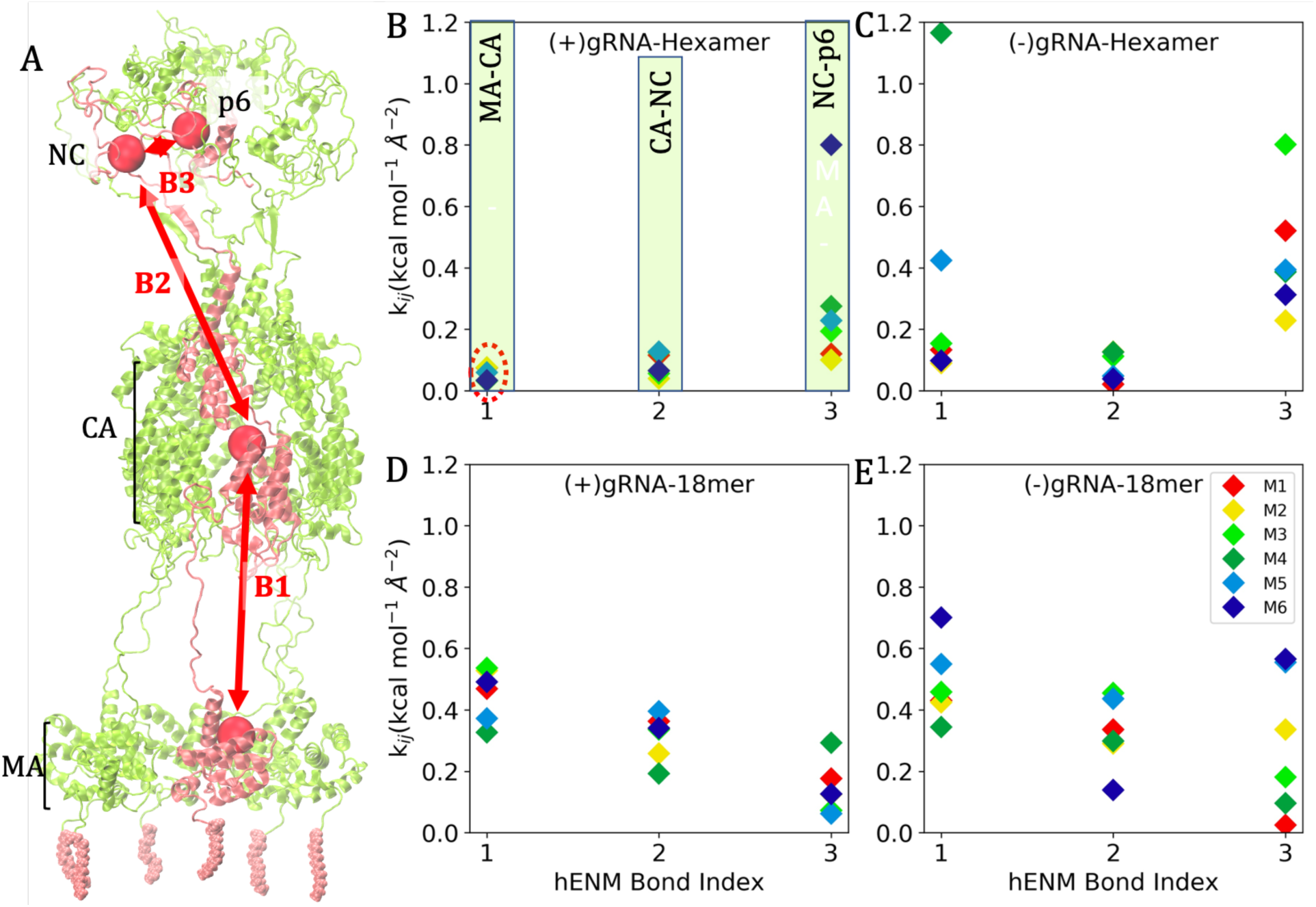
Elastic force constants (k_ij_) for hENM effective harmonic bonds formed between interacting Gag domains. A. The construction of the four-site CG model for the hexameric Gag model. (B-C) Dynamical correlation between domains (estimated as k_ij_) for six Gag monomers (M1-6) of the hexameric Gag models, with and without implicit gRNA binding. (D-E) k_ij_ values for six Gag monomers in the 18-mer Gag model constituting the inner hexameric ring. The hENM bond IDs 1, 2, and 3 correspond to MA-CA, CA-NC, and NC-p6 interactions, respectively.

In the following sections, we describe the simulations models, present the analyses of the structure and dynamics of Gag polyprotein at different multimerization states, as well as lipid sorting, and discuss our findings in the context of the current knowledge of HIV-1 assembly and Gag multimerization process.

## Results

### All-atom molecular dynamics (AAMD) simulations of Gag multimers with asymmetric membrane

We have carried out large-scale (∼ 4.5M atoms) AAMD simulations of Gag multimers bound to an asymmetric membrane. An atomic model of a full-length Gag was constructed using experimental cryo-ET and NMR-resolved structures of different domains. Our membrane model is based on lipidomic analysis of HIV-1 particles produced in HeLa cells reported by Lorizate *et al.* (*63*) (details of AA models are discussed in the ***Materials and Methods*** section). The inner leaflet of the bilayer contains anionic phospholipids such as phosphatidylinositol 4,5-bisphosphate (PIP2), and phosphatidylserine (PS) that promote membrane binding of the myristoylated MA (Myr-MA) domain, and the outer leaflet is enriched with raft-forming sphingomyelin (SM) and cholesterol. Pre-built models of Gag 18-mer, hexamer, and dimer were carefully equilibrated with the pre-equilibrated membrane patch where the MA domains were allowed to bind the inner leaflet of the asymmetric membrane (**Figure 1**). In the hexameric Gag bundle system, all six Myr groups were inserted into the membrane (***Figure 3***), whereas only the inner twelve Myr groups of the 18-mer Gag bundle were inside the membrane throughout the final trajectories (**Figure 1**). The IP6 anion maintains the stability of 6HB of the hexameric Gag bundle and the inner hexameric ring of 18-mer. Due to the lack of availability of an atomic model structure of gRNA bound to Gag multimers, in order to examine the general influence of gRNA binding on the structure and dynamics of Gag polyproteins, our modeling technique emulated the influence of gRNA binding on the dynamics of the NC domain through an effective or “implicit” interaction (see ***Materials and Methods* and Figure S1 of the Supporting Information**).

**Figure 3:**
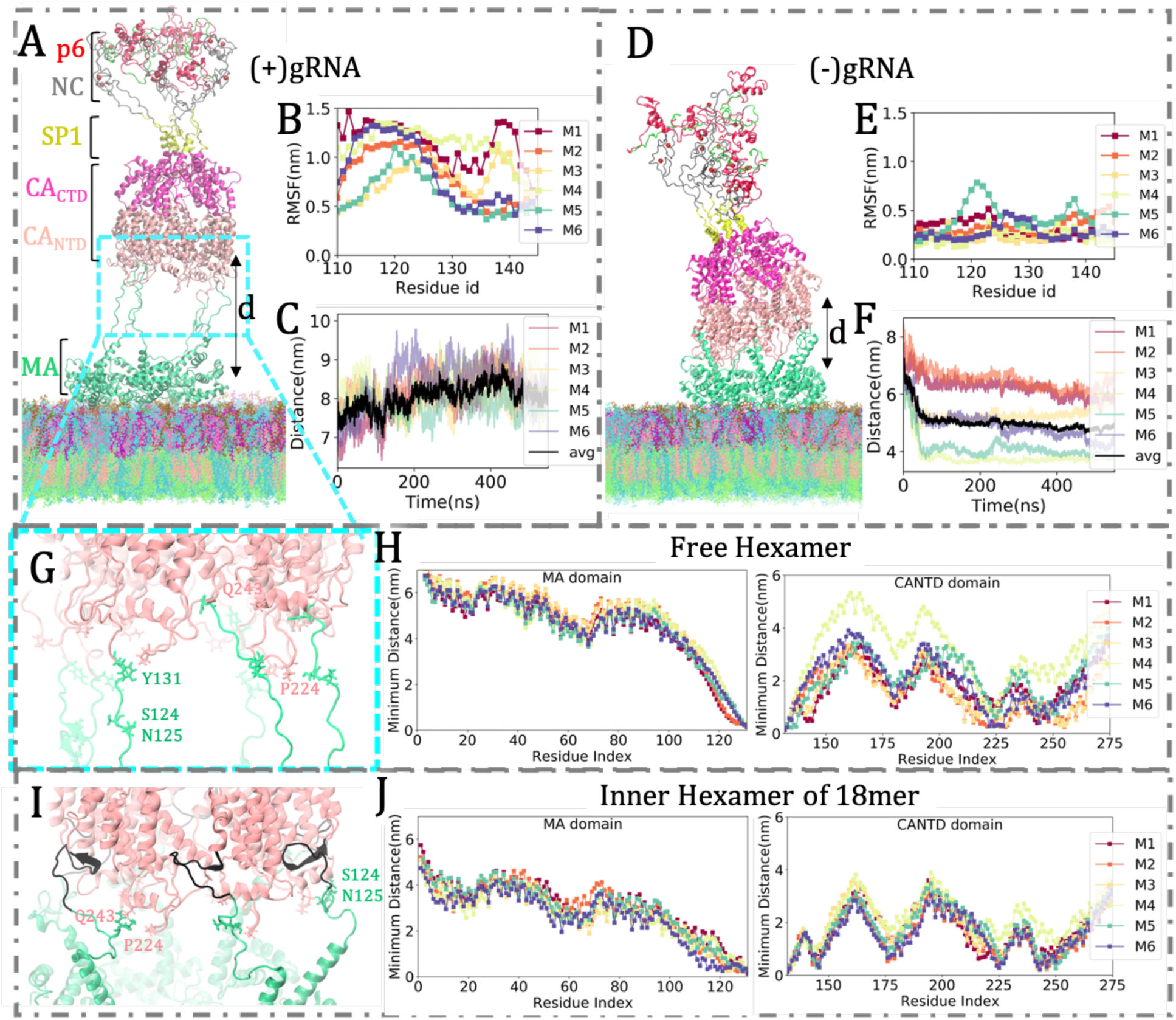
Structure and dynamics of MA-CA linker domain in hexameric Gag bundles. (A,D) Snapshots of the membrane-bound free hexameric Gag bundles in the presence and absence of implicit gRNA binding, respectively. (B,E) Root-mean-square-fluctuation (RMSF) of residues in the MA-CA linker region, (C,F) Center-of-mass (COM) distance between MA and CA_NTD_ domains of the six Gag monomers (chain M1-6) and the average distance between them. (G) Interaction between MA and CA_NTD_ domain in the presence of implicit gRNA binding (shown in (A)), (H) Minimum distance of residues of MA/CA_NTD_ domain from CA_NTD_/MA domain, respectively, (I-J) Same as (G-H), but for the inner hexamer of 18-mer Gag. Positions of MA residues, such as SER124, ASN125, and TYR131, are highlighted in green, and CA_NTD_ residues, such as PRO224 (CypA loop), GLN243 (helix5) are shown in pink color.

### Inter-domain correlation of Gag proteins is influenced by the multimerization state and (implicit) gRNA binding

We have leveraged CG modeling techniques as an analysis tool in order to gain an understanding of the effective interactions between MA, CA, NC, and p6 domains. Four-site CG models of Gag proteins for all the simulated systems were parametrized using the EDCG and hENM methods (see **Materials and Methods**). For this purpose, we have considered the free hexameric Gag model and the inner hexameric Gag bundle of the 18-mer system, in the presence/absence of implicit gRNA binding. Each main structural domain of Gag is represented by one CG group (i.e., “site” or “bead”), mapped by the EDCG method. Depending on the essential dynamical model of the domain, the CG bead may or may not be close to the center of mass of the domain. The effective harmonic force constants between CG beads, denoted as *k*_ij_ and determined by the hENM method, quantify the interaction strength between Gag domains.

Figure 2 presents k_ij_ values for hENM bonds B1, B2, and B3 corresponding to MA-CA, CA-NC, and NC-p6 interactions, respectively. The higher value of *k*_ij_ signifies a stronger correlation between the two domains. The implicit gRNA binding, denoted by (+)gRNA, is observed to impact not only CA-NC and NC-p6 but also MA-CA interactions in both the hexameric and 18-mer Gag model systems. The *k*_ij_ values for MA-CA interactions in the (+)gRNA-hexamer model are the lowest among all the models, suggesting uncorrelated dynamics of the two domains upon gRNA binding (Figure 2B). The MA-CA equilibrium separation (*x*_0_) values of all six monomers for this model are higher than other models, as shown in **Figure S2**. In the inner hexamer of the 18-mer Gag model, the influence of gRNA implicit binding on the MA-CA interaction is also present; however, it is not as pronounced for all the Gag monomers (Figure 2D-E), **Figure S2**(D-E)). Furthermore, the MA-CA correlation for the Gag proteins in the inner hexamer of 18-mer is observed to be higher than the free hexameric analog. Together, our findings reveal the dependence of interactions between neighboring domains on the multimerization state of Gag proteins and the implicit gRNA binding. The gRNA binding in the free hexameric model eliminates the dynamical correlation between MA and CA domains. By contrast, at the higher multimeric state ((+)gRNA-18 mer), the Gag polyprotein retains the MA-CA correlation, which has implications for the collective Gag bundle interaction with the membrane.

### Molecular interactions involving disordered MA-CA linker domain of Gag

Here we focus on the less explored domain of Gag multimers, the flexible linker between MA and CA domains, which contains a proteolytic cleavage site. Our simulated models allow us to study the structure and dynamics of the MA-CA linker in the inner hexamer of the 18-mer Gag lattice and compare that with the free hexamer, with/without implicit gRNA binding. The final structure of the free hexameric bundle structure bound to the asymmetric membrane with an implicit gRNA binding to its NC domain, root mean square fluctuation (RMSF) of the MA-CA linker domain residues, and the center-of-mass(COM) distance between MA and CA_NTD_ domains for all the six Gag monomers are shown in Figure 3A-C. The average distance between the COM of the hexameric CA and MA domains is shown in black. Evidently, MA-CA linkers in this hexameric bundle adopt an extended conformation, and most of the residues (starting from SER110) in the six linkers remain in a random-coil conformation. In contrast, the free hexameric bundle without an implicit gRNA binding to the NC domain (denoted by (–)gRNA) attained a conformation where the MA-CA linker domain of all Gag monomers are contracted allowing direct contacts between these two domains (Figure 3D-F). RMSF values of all the MA-CA linker residues of this model are considerably lower than the former (Figure 3B, E). The average COM distance between MA and CA in this system is ∼3 nm less than the former (Figure 3C, F). In the (–)gRNA hexameric bundle, our simulations sampled direct interactions between CA_NTD_ and MA domain residues. A hydrophobic collapse is observed in some of the monomers of this system, mediated by hydrophobic residues of helix2 of CA_NTD_, ILE146, ALA145, and C-terminus of the MA domain (ALA117, ALA118, ALA119).

A comparison of the conformations of this linker domain in the free hexameric bundle (Figure 3G, H) with the inner hexamer of the 18-mer Gag (Figure 3I, J) reveals altered characteristics of MA-CA linker domain induced by Gag multimerization. Here we compare two systems in the presence of implicit gRNA binding. In the inner hexamer of the 18-mer Gag, MA domains up to residue 115 (at least) remain in the helix form (Figure 3I). In the free hexamer, a portion of the linker domain (residues 120-130) binds CA_NTD_ domain residues transiently; however, these interactions are stable in the inner hexamer of 18-mer Gag. The minimum average distances of MA and CA_NTD_ domains are computed to quantify the interactions (Figure 3H, J). The structure of the CA_NTD_ domain consists of seven α-helices and CypA binding loop between helix4 and helix5 (shown in **Figure S6)**. This flexible loop is an extended proline-rich loop (residue 218-226) that mediates the binding of a host cell restriction factor cyclophilin A (CypA). CA_NTD_ residues which remain in close proximity to the MA domain belong to CypA binding loop and helix5 (residues 244-252). PRO224 and GLN243 of CA_NTD_ domains are highlighted in Figure 3G, I.

In addition, the N terminal residues of the CA_NTD_ domain in the free hexamer remain in a random coil conformation, whereas in the inner hexamer of 18-mer, MET141-PRO148 participate in CA_NTD_-CA_NTD_ inter-Gag dimeric interactions by forming antiparallel beta-sheet which is stabilized by hydrophobic residues such as VAL142, ALA145 and ILE146 (Figure 3I, **Figure S4**). The dynamics of this domain have been investigated in two types of hexameric systems. As depicted by the RMSF values, residues 141-148 in the inner hexamer of 18-mer are much more structured, and in turn, make the whole MA-CA linker considerably less flexible than the free hexameric Gag (**Figure S4**). An important point to note here is, as CA_NTD_ trimeric interface is anticipated to drive MA trimerization, the dimeric CA_NTD_ interactions can contribute to the Gag multimerization by promoting MA trimer-trimer interactions to form immature MA hexamer-of-trimers lattice structure. Based on the cryo-ET density map, Schur *et al.* have reported the dimeric interface between CA_NTD_ domains, mediated by helix1 of CA (22), here we report the contribution of the N-terminal loop domain of CA_NTD_ in stabilizing this dimeric interface. In summary, our findings here reveal the influence of gRNA binding and Gag multimerization on the structure and dynamics of the MA-CA linker domain. This linker domain remains in a collapsed state in the free hexameric Gag bundle without gRNA binding. With gRNA binding in the free hexameric state, this linker domain exhibits an extended disordered state where a portion of the linker domain (residues 120-130) binds CA_NTD_ domain residues transiently. However, these interactions become stable in the 18mer state and additional interactions mediated by MET141-PRO148 stabilize the dimeric interface between two Gag molecules.

### Implicit gRNA binding influences the dynamic organization of the membrane

Interestingly, our AAMD simulations with an asymmetric membrane revealed specific and non-specific MA-lipid interactions and lipid sorting by Gag multimers. Figure 4A shows the interactions of a MA monomer (part of the hexameric Gag bundle) with PIP2 and PS lipids when the Myr group is inserted into the membrane, in a stable configuration. A flexible loop at the membrane binding interface of MA protein enriched with basic amino acid residues ARG and LYS (residues 17-31), is denoted as the “highly basic region” (HBR) (64). Membrane targeting of Gag is driven by the specific interactions between the HBR domain and PIP2 lipids, key residues participating in these interactions being ARG21, LYS26, LYS29, and LYS31 (as shown in Figure 4A)(17,65). Apart from these, other PIP2 binding sites include HBR residues (ARG19, LYS25), N terminal residue (ARG3), and helix2 residue (ARG38). The membrane binding mechanism also includes (i) non-specific interactions of basic residues of MA at the membrane binding surface with monovalent anionic phospholipid, phosphatidylserine (PS) (Figure 4A) and (ii) hydrophobic interactions of Myr residues with the lipid tails.

**Figure 4:**
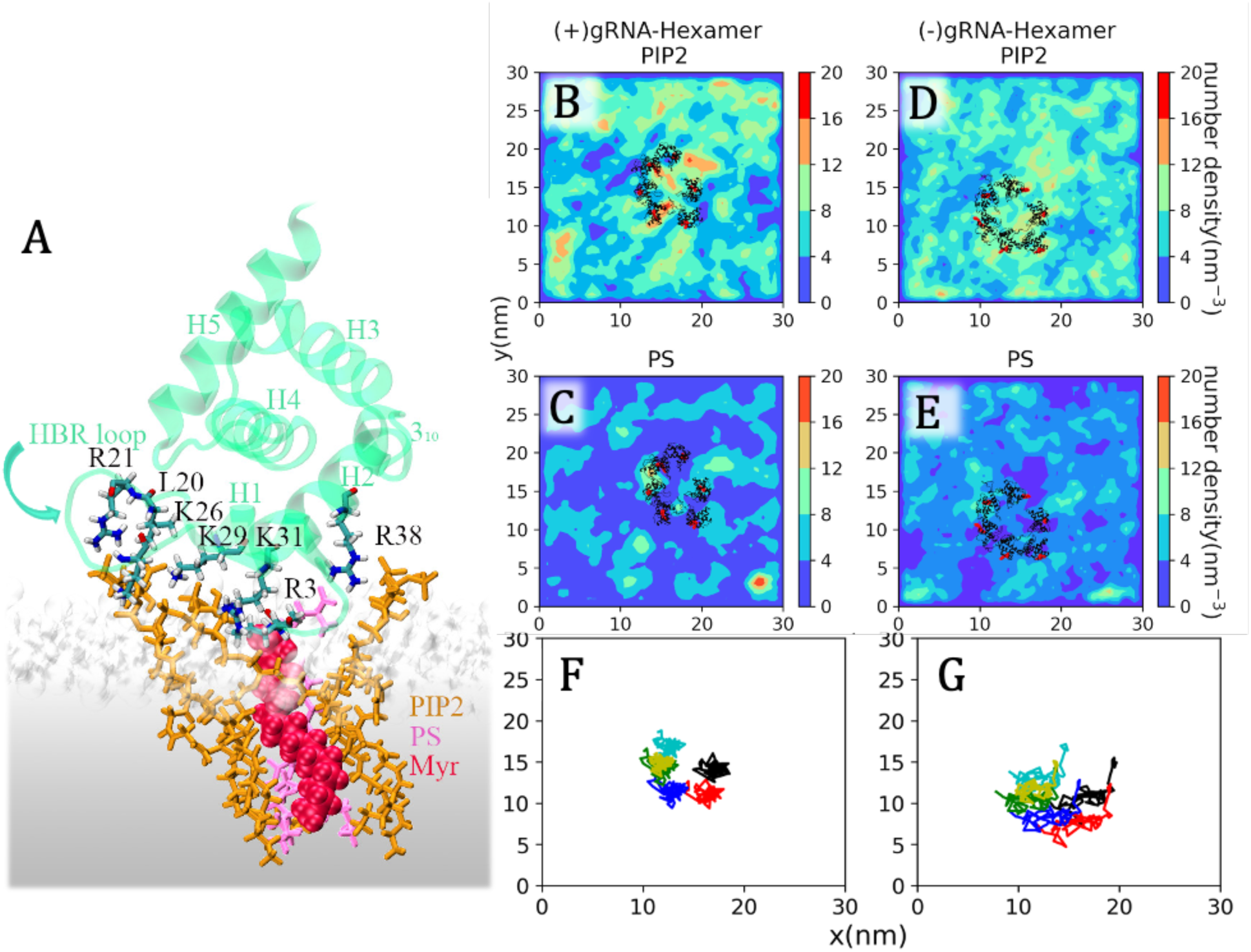
MA-membrane interactions in Gag multimers. (A) Membrane-bound matrix (MA) monomer, belonging to the hexameric Gag bundle system. Amino acid residues participating in specific/non-specific binding with PIP2 and PS lipids are shown. (B-E) Time-averaged PIP2 and PS density maps around six separated Myr-MA monomers of hexameric Gag bundle, with implicit gRNA binding(B,C), without gRNA binding to NC domain (D,E). The average positions of MA and Myr are shown in black and red, respectively. (F,G) Trajectories of the center-of-mass of the MA domain during the timeframes used to compute time-averaged density maps of lipids. The reduced lateral mobility of the MA domain in (+)gRNA-hexamer results in the dynamical localization of lipid molecules.

We also investigated lipid arrangement around the MA domain of the hexameric and 18-mer Gag bundles. The number density of the headgroup atoms of PIP2 and PS lipids are computed with the last 500 ns of the simulated trajectories. Figure 4B-E present the time-averaged lipid density maps of PIP2 and PS at the inner leaflet of the bilayer surrounding six separated MA proteins, belonging to the hexameric Gag bundle with/without implicit gRNA binding. The higher density domains in the time-averaged density maps of (+)gRNA hexamer (Figure 4B,C) can be caused by either an enrichment of such lipid molecules in the vicinity of the MA domain or the dynamic localization of lipid molecules. The simulated trajectories did not show a significant difference in the number of MA-bound PIP2/PS lipids in these two systems. However, the lateral mobility of the MA domain of the (+)gRNA hexameric Gag bundle was observed to be reduced compared to the (–)gRNA hexameric Gag bundle (Figure 4F,G) which modifies the dynamic organization of the MA-bound lipid headgroups. The dependence of the lateral dynamics of membrane-bound Gag molecules on the gRNA binding, as captured in our simulations, is in agreement with prior experimental data(43). These density maps were reproduced with different trajectories of both systems.

We further investigated the lipid sorting behavior of the 18-mer Gag bundle. The specific MA-lipid interactions depend on the exposed surface electrostatic potential and, in turn, the oligomerization state of MA domains. The lipid density maps around the 18-mer MA domain exhibit a higher density of PIP2 and PS in the hole region as the immature MA hexamer-of-trimer structure is comprised of an inner ring of six HBR domains (**Figure S5**).

### gRNA binding modulates the binding surface of 18-mer Gag

Our simulations reveal that gRNA binding (treated implicitly, see Methods) mainly modulates the structure and dynamics of the SP1 domain of the 18-mer Gag bundle which plays a crucial role in the multimerization process. In both the 18-mer Gag models, shown in Figure 5A-B, IP6 binds to the inner six-helix bundles (6HB) (37), not the outer twelve Gag CASP1 domains. We have

**Figure 5:**
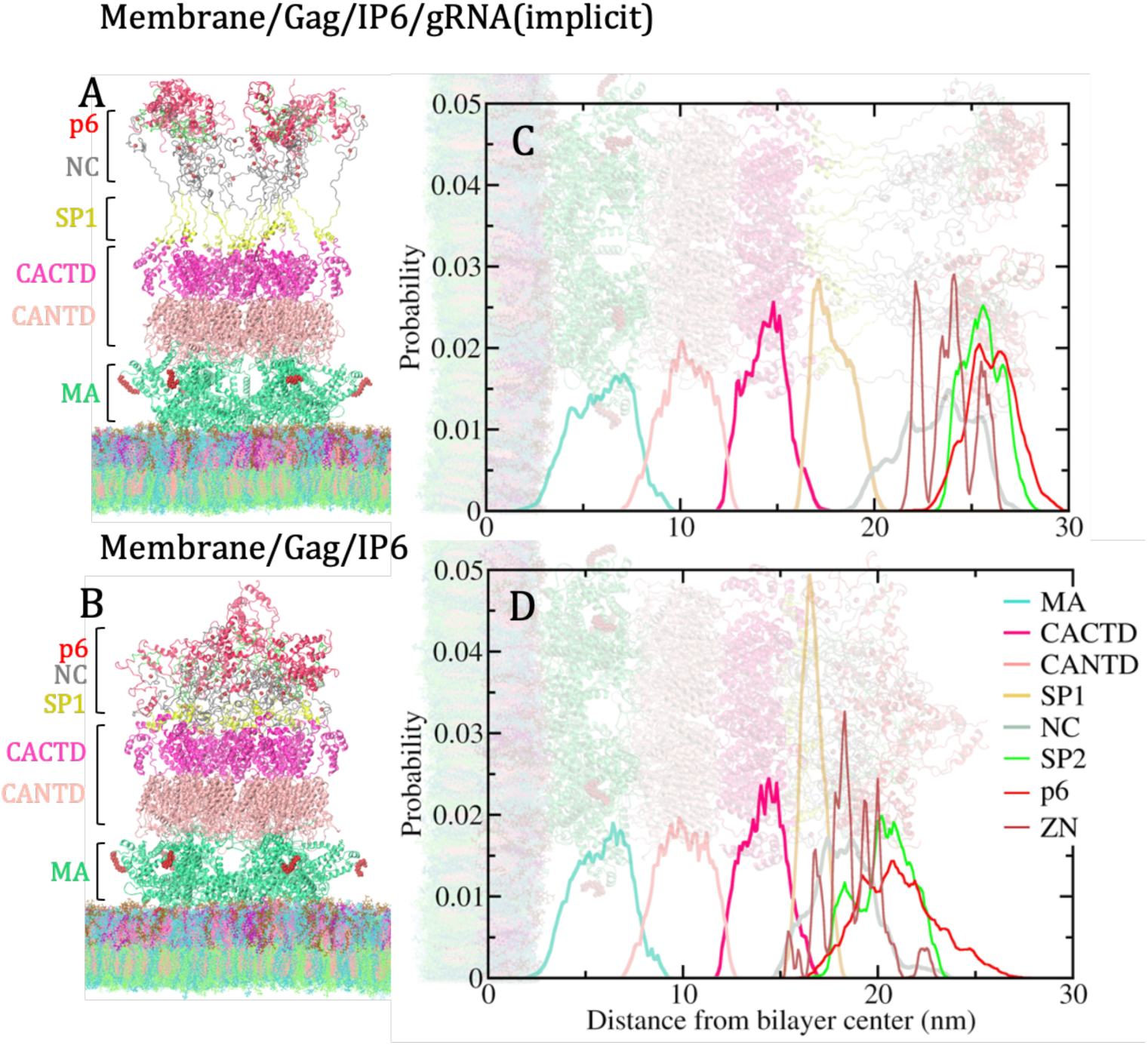
Snapshots of membrane-bound Gag 18-mer. Final structures of microsecond-long simulations of 18-mer Gag (A) with an implicit gRNA binding, (B) without gRNA binding, (C-D) Probability density of the Cα atoms in the different domains of the outer twelve monomers of the 18-mer Gag bundles along the membrane normal direction. Without gRNA binding, the SP1 domain of outer monomers of 18mer Gag, being buried inside other domains, disrupts an efficient binding interface.

computed the probability density of the Cα atoms in the different domains of the outer twelve Gag monomers using our AAMD simulation trajectories and plotted it against the z-distance from the bilayer center (along the membrane normal direction) (Figure 5C,D). Although there are minor differences in the arrangement of CA_CTD_ and CA_NTD_ domains, the SP1 helices of the outer monomers (shown in yellow color) are observed to be buried inside other domains in the absence of gRNA binding. The NC domain and CA_CTD_ domain interact with each other in this system. In contrast, in the presence of gRNA binding, SP1 domains keep CA_CTD_ and NC domains separated from each other. It is now generally accepted that SP1 helices play a predominant role in Gag multimerization through the formation of a six-helix bundle (6HB), stabilized by IP6 (37). In the next section, we present a detailed analysis of the altered characteristics of the SP1 domain.

### Conformational dynamics of SP1 helices

As mentioned earlier, the CA_CTD_-SP1 domain offers the predominant Gag-Gag binding interface and plays a crucial role in the recruitment of cytosolic Gag molecules as well as in the proteolytic cleavage process during viral maturation. We have analyzed the structure and dynamics of this domain in the outer twelve Gag monomers of the 18-mer Gag bundle (Figure 6A,E). This domain of the six pairs of outer Gag monomers forms partial hexamers, interacting with each other via the MHR loop and helix10 (Figure 1). The SP1 domains of these partial hexamers maintain stable binding while undergoing spontaneous helix-loop transition, partially. The helicity and solvent accessibility of the SP1 loop domain was examined, in detail. The solvent-accessible surface area (SASA) of the SP1 domain (residues 362-376) for all the 12 outer Gag monomers are higher and the fluctuation of SASA over time is also higher in the presence of implicit gRNA binding (Figure 6B-C). For any protein association process, solvent accessibility of oligomer forming protein domain is crucial(66). In this system, solvent accessibility of this SP1 domain might facilitate the efficiency to bind new Gag molecules and the growth of immature lattice structures. The role of dynamic helix-coil transition of this domain during viral assembly and maturation has also attracted attention in the past. Experiments revealed that these SP1 helices act as a “molecular switch” for Gag multimerization (31,67,68). Figure 6D shows that up to 500 ns of the trajectory, helicity fluctuates significantly and thereafter converged to a value of ≤ 3, for most of the helices. On the other hand, in the absence of gRNA binding to the NC domain, helicity ranges from 2-6 for different SP1 helices without much fluctuation. The calculation of helicity was performed using PLUMED(69) and the magnitude of helicity in Figure 6D,H signifies the number of six amino-acid residue segments that are in an alpha-helical configuration. All these results, together, signify that the gRNA binding induces a change in environmental condition, i.e., molecular crowding and polarity of the medium around this domain, and modulates SP1 conformation.

**Figure 6:**
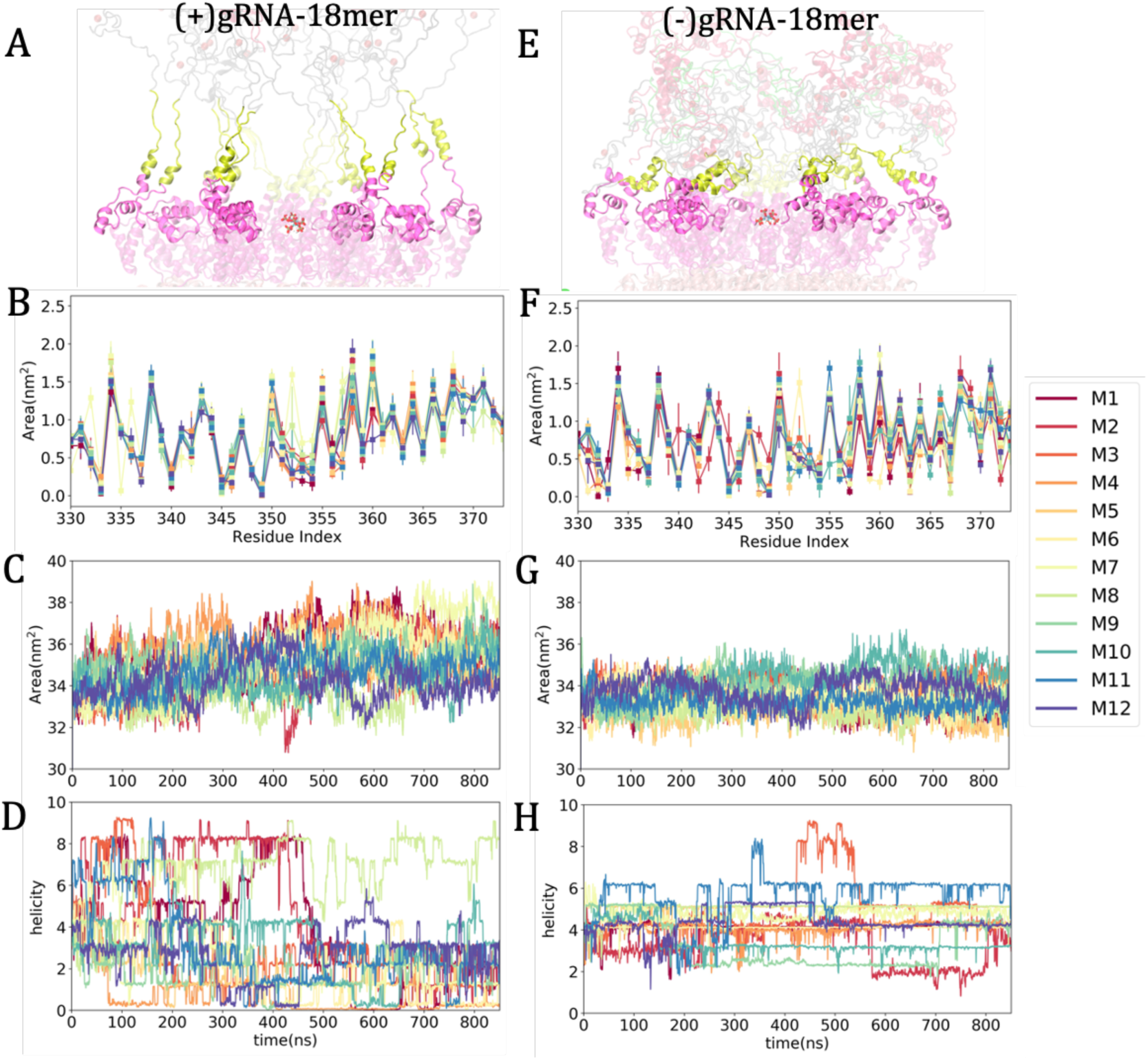
Characterization of the CA_CTD_-SP1 binding surface. (A, B) CA_CTD_-SP1 domain of the outer twelve monomers of 18-mer with/without gRNA binding. CA_CTD_ and SP1 domains are shown in pink and yellow, respectively. (B, F) Solvent accessible surface area (SASA) of CA_CTD_-SP1 residues. (C, G) Time evolution of the total solvent accessible surface area of residues 340-372. (D, H) The helicity of the SP1 domain (residues 362-376) as a function of time. (A-D) in the presence of implicit gRNA binding, (E-H) without gRNA binding. gRNA binding is observed to increase the solvent accessibility of SP1 helices and induces partial helix-to-coil transition.

### Membrane-bound dimeric Gag-bundle collapsed in the absence of implicit gRNA binding

Unlike hexameric and 18-mer Gag bundles, simulated structures of membrane-bound dimeric Gag proteins exhibit a dramatic change in the presence and absence of the gRNA binding (Figure 7). Similar to the other two systems, here also we started our simulations from an extended state of the Gag proteins. Gag dimer ((+)-gRNA) exhibits an elongated and flexible MA-CA linker domain, conferring mobility to the MA domain on the membrane surface. The H1 helix of the CA_NTD_ domain also undergoes a partial unfolding. MA-CA linker domain in this system does not interact with any CA_NTD_ residues, as reported for the inner hexamer of Gag 18-mer and free hexameric Gag bundle (Figure 3 and **Figure S7**). Only the H1 helix of CA_NTD_ and the H9 helix of CA_CTD_ mediate Gag-Gag interaction in this system. Although the starting structure contains SP1 helices (as constructed from experimental data (37)), the simulation of (+)gRNA dimer leads to an unfolded coiled state of SP1. There was no IP6 anion bound to the SP1 domain in our simulated models of dimeric Gag. On the other hand, in the (–)gRNA Gag dimeric system, the Gag molecules become collapsed and almost all the domains of Gag interact with each other. Our simulated trajectory suggests that the independent movement of the NC domain in the (–) gRNA dimeric system allows them to interact with CA domain residues and subsequently other interdomain interactions result in this collapsed state.

**Figure 7:**
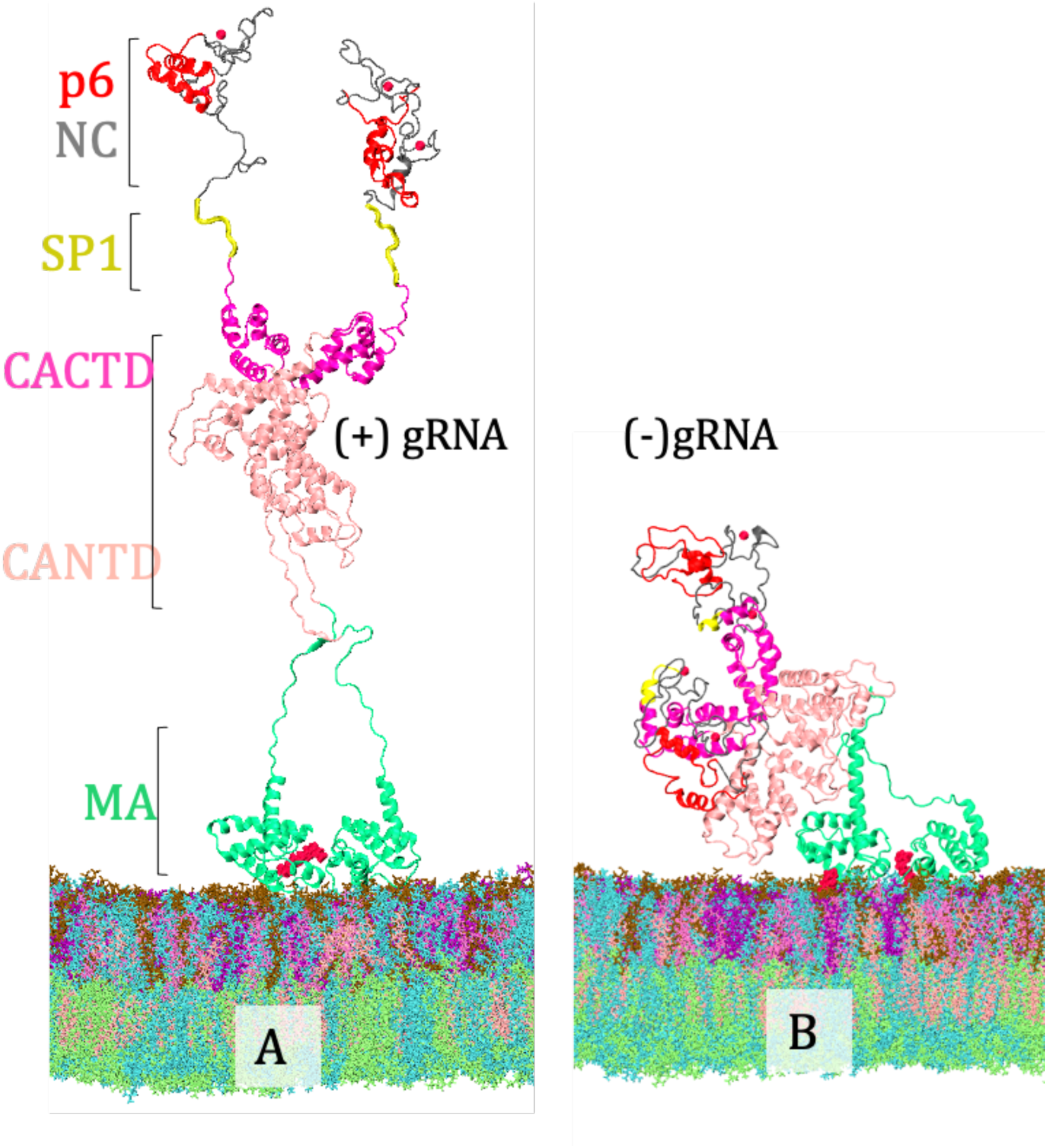
Membrane-bound dimeric Gag bundles. A. An extended form of Gag dimer in the presence of an implicit gRNA binding. B. Collapsed state of Gag dimer in the absence of gRNA binding.

## Discussion

Using large-scale AAMD simulations, the present study explores the behavior of Gag polyproteins in different multimerization states and the impact of a gRNA binding on the structure and dynamics of Gag. Being a multidomain protein, cooperative homo-domain interactions play a crucial role in the Gag multimerization process. At first, we investigated the inter-domain correlation of Gag in hexamer and 18-mer with a simple four-site coarse-grained model of Gag, where each structural domain of Gag has been represented by a CG site or bead. Using the AAMD trajectories, CG mapping was performed using the EDCG method, and the effective interactions between different CG sites are evaluated using the hENM method. As measured by the resulting k_ij_ values (Figure 2), MA-CA inter-domain interactions were found to depend on both the multimerization state of the Gag protein and the implicit gRNA binding. The simulated trajectories of the membrane-bound free hexameric Gag model ((+)gRNA) exhibit uncorrelated dynamics between MA and CA However, at the higher multimeric state ((+)gRNA-18 mer), the dynamic correlation between MA and CA domains is recovered.

Next, we studied the molecular-level interactions leading to the altered characteristics of the Gag molecules themselves. The presence/absence of gRNA binding to the NC domain is observed to modulate mainly the MA-CA linker domain of the hexameric Gag and the SP1 domain of the 18-mer Gag. The multimerization state of Gag also influences the dynamics of the MA-CA linker domain as it is observed to be less disordered in the inner hexamer of the 18-mer Gag bundle compared to the free hexamer (**Figure S4**). With gRNA binding, MA-CA linkers in the free hexameric Gag adopt an extended conformation, and residues 120-130 interact with CA_NTD_ domain residues transiently. In contrast to this, without gRNA binding the NC domain attained a conformation where the MA-CA linker domains of all Gag monomers are contracted, i.e., the COM distance between CA and MA domains is ∼3nm less than the former ((+)gRNA hexamer) and the C terminus residues of MA domain exhibit stable binding with CA_NTD_ residues, especially mediated by hydrophobic residues (Figure 3D-F). Also, in the inner hexamer of the 18-mer Gag bundle, we identified stable and transient interactions between the CypA binding loop and MA-CA linker, driving inter-domain, as well as inter-Gag, interactions (**Figure S7**). In the 18-mer Gag model, without a gRNA binding, the SP1 helices are observed to be buried inside other domains with reduced solvent accessibility (Figure 5 and Figure 6). Rein and coworkers have revealed that the SP1 conformation is sensitive to environmental conditions, including molecular crowding and polarity of the medium (68). Our findings agree with this experimental insight, as in the presence of gRNA binding the outer twelve SP1 helices in the 18-mer Gag become solvent exposed and exhibit a spontaneous helix-to-coil transition. On the other hand, in the dimeric Gag bundle, the SP1 helices remain solvent exposed and an unfolding of the SP1 helices is observed in the presence of gRNA binding. Our prior study showed that dimeric Gag is recruited to the immature growing lattice (32). Our present simulations suggest that (i) gRNA binding is crucial for dimeric Gag to have an efficient binding interface and (ii) the SP1 domain remains in a coiled state in (+)gRNA-dimeric Gag molecules. The coil-to-helix transition of the SP1 domain might require prior hexamer formation by the CA_CTD_ domain and the presence of IP6 anions.

The MA lattice structure in the immature HIV-1 particle has been recently revealed by cryo-ET experiment, in which Myr-MA trimers interact with each other with the N-terminal residues, H1 and 3_10_ helices (25). Several studies revealed that Gag assembly occurs within specialized microdomains of the plasma membranes enriched in anionic phospholipids, especially PIP2, and cholesterol (45). Gag multimerization is also observed to play a key role in PIP2 clustering (46). In our simulations of the Gag 18-mer models (with and without gRNA binding), a high density of PIP2 and PS is observed close to the inner hole of the hexamer-of-trimer (**Figure S5**). We computed time-averaged lipid density maps for hexameric Gag systems from the last 500 ns of the AAMD trajectories. Interestingly, the dynamic organization of PIP2 and PS headgroup in the (+)gRNA and (–)gRNA systems are observed to be different, the former being more dynamically localized than the latter (Figure 4B,C). This is induced by the reduced lateral mobility of the MA domains (Figure 4F).

As our Gag model contains all the domains including the 52-amino acid domain in the C-terminal of Gag, known as p6, we could investigate the less-explored interactions between the p6 domains. This domain is predicted to be mostly disordered and mediates the recruitment of the ESCRT complex that drives the viral budding (70). In addition, it was suggested that p6 domain plays a role in the specificity of the Gag-gRNA interaction (71). Under the membrane-bound condition, p6 is shown to adopt a two-helix structure in which shorter helix-1 is connected to a pronounced helix-2 by a flexible hinge region as shown by Solbak *et al.* (72). The N-terminal helix of p6 in some Gag monomers undergoes a helix-to-coil transition in the simulated trajectories of the (+)gRNA 18-mer Gag. Our simulations also explored stable p6-p6 interactions. The C-terminal helix of one Gag monomer binds to both the helices of another Gag chain (**Figure S8**). This kind of binding is stabilized both by hydrophobic and electrostatic interactions. The two Gag monomers interacting through the p6 domain do not involve in any other homo-domain interactions in our simulations.

Large-scale AAMD of Gag bundles in different multimerization states may encounter challenges with the timescale of Myr insertion. In the simulations of 18-mer Gag bundles ((+)/(–) gRNA binding), 12 out of 18 Myr groups are inserted into the membrane; however, in the hexameric Gag bundle, all six Myr groups are inserted into the membrane. On the other hand, within the simulated AAMD timescale, none of the Myr groups in the (–)-gRNA Gag dimer and only one Myr group in the (+)-gRNA Gag dimer system become fully inserted into the membrane. The other Myr group remained on the protein-membrane interface in an exposed state and kept “searching” for a large-enough lipid packing defect suitable for insertion. Our simulations, therefore, suggest that for the higher-order Gag multimers, the presence of a greater number of Myr-MA domains might enhance the lipid packing defects for Myr insertion.

An important point to note about our AAMD simulations is that there is so far no atomic model of Gag-bound gRNA is available to be used in such a simulation study. To examine the influence of gRNA binding on the structure and dynamics of Gag polyproteins, our modeling technique, therefore, sought to emulate the effect of the NC-gRNA binding on the dynamics of the NC domain. The weak restrained potential applied on the relative distance of the ZN atoms of the ZN-finger domain only hinders the independent movement of the NC domain and does not fix any atoms/domains in space. Furthermore, the study of protein association/dissociation remains a major challenge for computational studies (66,73,74). For a multidomain protein like Gag, the direct AAMD simulation study of the multimerization process is prohibitively difficult. In the present work, we thus aimed to perform a molecular-level characterization of the structure and dynamics of the Gag protein in different states. In the future, the study of long-time and large-scale processes like the structural transition of membrane-bound Gag during the viral assembly process will be addressed by multiscale, coarse-grained simulation methods.

To summarize this work, using different model systems of Gag dimer, hexamer, and 18-mer, we investigated via AAMD (i) how the characteristics of Gag change during the multimerization process, and (ii) the influence of the independent or restrained movement of the NC domain (caused by gRNA binding) on the interdomain dynamic correlation of Gag. In the hexameric Gag bundle, the conformational dynamics of the MA-CA linker domain and its interaction with the CA_NTD_ domain, especially CypA binding loop and helix5 residues, become altered upon multimerization of Gag and gRNA binding. In the 18-mer Gag, the solvent accessibility of SP1 helices changes without gRNA binding, which modulates the binding surface and may further impact the recruitment of cytosolic Gag molecules. The independent motion of the NC domain in the dimeric Gag bundle (in the absence of gRNA) results in a collapsed state of the dimer where almost all the domains directly interact with each other. Furthermore, in the hexameric Gag bundle, along with the alteration of the structure and dynamics of the MA-CA linker domain, the gRNA binding also facilitates the localization of the PIP2 and PS lipid headgroups around the Myr-MA domain. Future work will be necessary to better unravel the cooperativity between homo-domain multimerization of Gag and gRNA packaging by a growing Gag lattice. We intend to systematically determine coarse-grained (CG) inter-protein interactions parametrized directly from the all-atom (AA) configurations from the 18-mer Gag bundle simulations by utilizing a systematic “bottom-up” coarse-graining methodology (75) and then explore how Gag-Gag, Gag-gRNA, and Gag-membrane interactions coordinate the viral assembly processes, by using these CG simulations.

## Materials and Methods

### Simulation setup for different Gag multimers

The initial atomic model of full-length Gag was constructed using a fitted model structure derived from the experimental cryo-ET density map of immature MA hexamer of trimers (EMD-13087) (25), CASP1 oligomers (PDB ID: 5L93, 6BHR) (37), NMR structure of NC domain with CCHC ZN finger motifs (PDB ID: 1A1T) (26,27) and p6 domain (PDB ID: 2C55) (28). MA trimers (PDB ID: 1HIW) were fitted into the cryo-ET density map and the monomeric MA structure (PDB ID: 1UPH) with all MA residues upto TYR131 was then superimposed to MA assemblies to obtain the final MA assembly structure (11,15). Myr tails, in an exposed conformation, were covalently attached to its N-terminal GLY1 residue using the CHARMM-GUI input generator (76). A fully hydrated lipid bilayer was built and equilibrated using CHARMM-GUI Membrane Builder and Quick-Solvator, respectively (76–80). Next, protein coordinates were positioned close to the inner leaflet of the membrane using VMD software (81), and those lipid molecules, that overlapped with Myr groups, were removed. The protein-membrane merged structures for the dimer, hexamer and 18-mer Gag bundles were solvated in 150 mM aqueous KCl solution with initial box lengths of 30 nm × 30 nm × 48 nm to allow enough water layers to be present between the protein units and the periodic image of the outer leaflet. The total size of both systems was ∼ 4.5M atoms.

### Asymmetric Membrane model

The membrane model designed for this study was based on lipidomic analysis of HIV-1 particles produced in HeLa cells reported by Lorizate *et al.* (21,63). The approximate composition of the extracellular leaflet was 35% cholesterol, 30% phosphatidylcholine (PC), and 35% sphingomyelin (SM) and the cytoplasmic leaflet was 15% cholesterol, 40% PC, 15% PE, 15% phosphatidylserine (PS) and 15% phosphatidylinositol (PIP2). All of the data shown in this manuscript were computed for the membrane system with DOPC, POPE, DOPS, SAPI, LSM, and cholesterol. However, the specific MA-lipid interactions have been verified using another membrane system with POPC, POPE, POPS, SAPI, LSM, and cholesterol. Our membrane model mimics the asymmetric nature of the HIV-1 membrane. A bilayer with a surface area of 30 nm x 30 nm was built using CHARMM-GUI Membrane Builder and Quick-Solvator (76–80) following a multistep minimization and equilibration protocol. This system was further equilibrated for 500ns before merging the equilibrated protein coordinates using VMD software (81).

### Molecular Dynamics (MD) Simulations and Analysis

We have considered three model systems for the AAMD simulations, i.e., Gag 18-mer, hexamer, and dimer, and simulated those with an asymmetric membrane model (see above subsection). All simulations used the CHARMM36m force field (82) and were performed in GROMACS 2019

(83). Energy minimization was performed using the steepest descent algorithm until the maximum force was less than 1,000 kJ mol^−1^nm^−1^. Then, equilibration was performed with harmonic restraints (using a 1,000 kJ mol^−1^nm^−2^ spring constant) on each heavy atom throughout the protein for 1 ns in the constant NVT ensemble with a time step of 1 fs, followed by a 15 ns constant NVT MD run with a time step of 2 fs and then a 15 ns MD run in the constant NPT ensemble with a time step of 2 fs. Next, replicas of all systems were simulated for 500 ns, respectively, for both systems in the constant NPT ensemble, following a parallel cascade selection MD procedure (84), to allow Myr insertion to occur. During this phase, Gag assembly structures were kept intact by applying harmonic restraints (1000 kJ mol^−1^nm^−2^) on Cα-backbone atoms of protein monomers, except for the first ten N-terminal residues of MA including the Myr group, as Myr insertion requires conformational flexibility of those residues. Within the simulation timescale, all 6 Myr groups of Gag hexamer, 12 out of 18 Myr of Gag 18-mer, and one of the two Myr groups (for the (+)gRNA model) were inserted into the inner leaflet of the bilayer. Next, multiple 1 μs trajectories were generated for all the Gag 18-mer, hexamer, and dimer systems. Prior restraints on the protein backbones were removed for the AAMD production runs. For these immature Gag multimers, to emulate the effect of gRNA binding to the NC domain, we have applied weak harmonic restraints (50 kJ mol^−1^nm^−2^) to the ZN atoms in the ZN-finger motifs of the NC domain to maintain the relative distance of ZN atoms while moving freely in water (**Figure S1**).

Throughout this AAMD procedure, the temperature was kept constant at 310.15 K using the Nosé –Hoover thermostat with a 1.0 ps coupling constant (85,86), and the pressure was set at 1 bar and controlled using the Parrinello-Rahman barostat semi-isotropically due to the presence of membrane, the compressibility factor was set at 4.5 ×10^-5^ bar^-1^ with a coupling time constant of 5.0 ps (87,88). Van der Waals interactions were computed using a force-switching function between 1.0 and 1.2 nm, while long-range electrostatics were evaluated using Particle Mesh Ewald (89) with a cutoff of 1.2 nm and chemical bonds containing hydrogens were constrained using the LINCS algorithm (90).

The linker domains of the atomic model of Gag were constructed using Modeller version 10.0 (91). The MA model fitting to the cryo-ET density map was done in ChimeraX (92). Protein-protein and protein-lipid interactions were visualized using VMD 1.9.3. (81). Analyses of the trajectories were performed using GROMACS 2019(83), PLUMED(69), MDAnalysis (93,94), and VMD Tcl scripts(81).

### Essential Dynamics Coarse Graining (EDCG)

The EDCG method (61), was used to define a CG model of Gag where CG sites correspond to four main structural domains of Gag polyprotein (MA, CA, NC, p6), (see Figure 2). In combination with principal component analysis (PCA), this method preserves the key essential dynamics of the system by maximizing the overlap of the CG model motions with the AA collective motions. The Cα atoms are grouped into CG sites by minimizing the following variational residual:

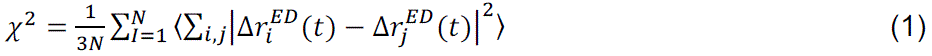

where *N* is the number of CG sites. The contribution of the displacement of residue i at time t to the essential subspace (spanned by essential modes) is denoted by 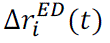. If the same quantity for residue j is of similar magnitude, they are identified as dynamically correlated and they become part of the same CG sites. In this study to build a four-site CG model of Gag, the displacement differences of the residues were evaluated using the last 400 ns of the AAMD trajectory when the fluctuations of Cα displacements were around steady values.

### Heterogeneous Elastic Network Model (hENM)

We parametrized hENM models for each simulated system to understand the effective inter-domain correlation in the Gag monomers being a part of the hexameric Gag model and the inner hexamer of the 18-mer Gag model (62). Each structural domain of Gag (MA, CA, NC, and p6) was represented by one CG bead, mapped by the EDCG method (Figure 2). The last 200 ns of trajectories were used to compute effective pairwise harmonic interactions (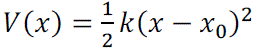) between CG sites, within a distance cut-off range. In this method, starting from a simple ENM (elastic network model) (95,96) with equal force constant for all the elastic bonds, k_ij_ is updated iteratively using the following equation, until the fluctuation in the CG model converges to the atomic data:

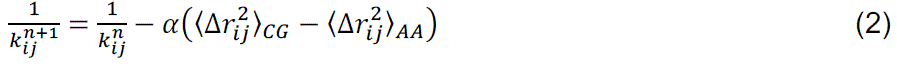

where 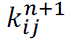 is the harmonic force constant for CG bead pairs i,j at iteration 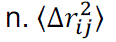 = 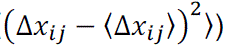 is the mean-squared distance fluctuation for i,j CG pairs and α is the scaling factor that controls the magnitude of the adjustment for each iteration. We have chosen α to be 0.5 and a cut-off distance of 10.5 nm for hENM calculations of all simulation systems.

## Author Contributions

P.B. and G.A.V. designed the research. P.B. performed simulations, analyzed the data, prepared the figures, and wrote the manuscript. G.A.V. supervised the research, writing process, contributed to discussions, edited and finalized the manuscript.

## Declaration of interests

The authors declare no competing interests.

## Acknowledgments

This research was supported by the National Institute of Allergy and Infectious Diseases (NIAID) for the Behavior of HIV in Viral Environments (B-HIVE) Center of the National Institutes of Health (NIH grant U54 AI170855). Computational resources were provided by the University of Chicago Research Computing Center (RCC), the Frontera supercomputer at the Texas Advanced Computer Center funded by the National Science Foundation (OAC-1818253), the Stampede2 supercomputer at the Texas Advanced Computing Center (TACC), the Extreme Science and Engineering Discovery Environment (XSEDE) supported by NSF grant ACI-1548562. (prior to September 2022), the Advanced Cyberinfrastructure Coordination Ecosystem: Services & Support (ACCESS) program (after September 2022) supported by NSF grants numbers 2138259, 2138286, 2138307, 2137603, and 2138296.

## Notes

### Competing Interest Statement

The authors have declared no competing interest.

