## Supporting Information for "Conformational transitions of the HIV-1 Gag polyprotein upon multimerization and gRNA binding"

**Affiliations:**

Department of Chemistry

The University of Chicago

5735 S. Ellis Ave, SCL 123

Chicago, IL 60637

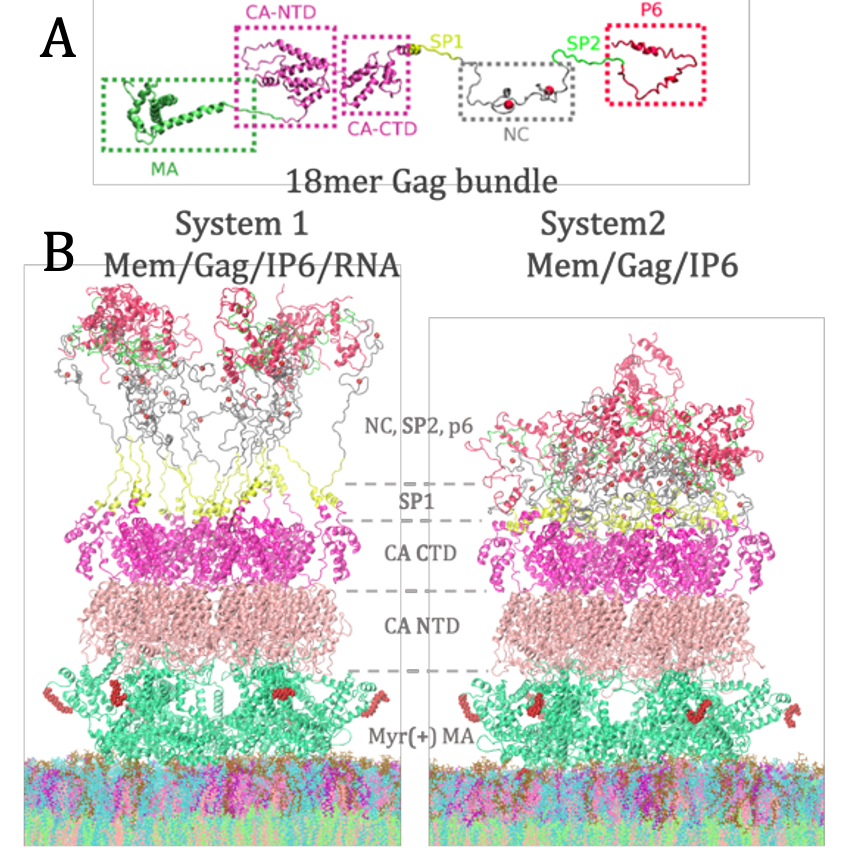

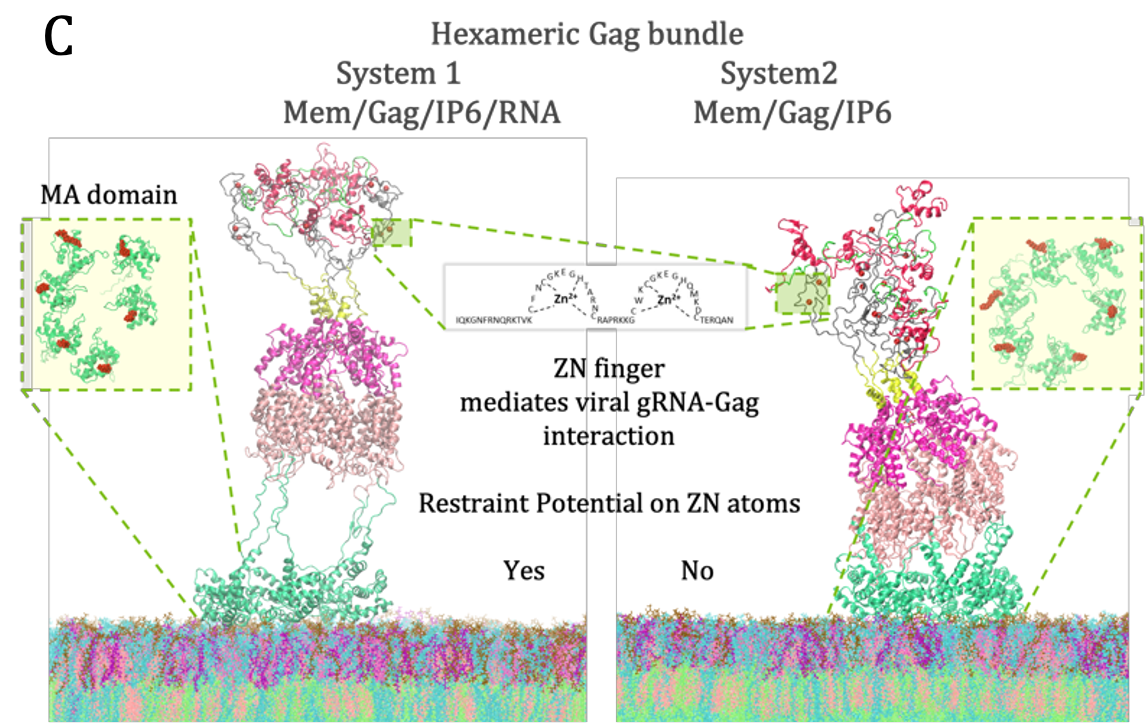

**Figure S1: Model systems for all-atom molecular dynamics simulations.** A. An Atomic model of the full-length Gag was constructed using experimental cryo-ET, X-ray, and NMR spectroscopy-derived structures of different domains. B. 18mer full-length Gag bundle bound to an asymmetric membrane with 12 out of 18 Myr group, inserted into the membrane, C. Hexameric Gag bundle bound to an asymmetric membrane where all 6 Myr groups anchor hydrophobic domain of inner leaflet. The binding of gRNA to the NC domain is emulated by applying a harmonic restrained potential to the ZN atoms of ZN-finger motifs in System1 of both Gag multimers.

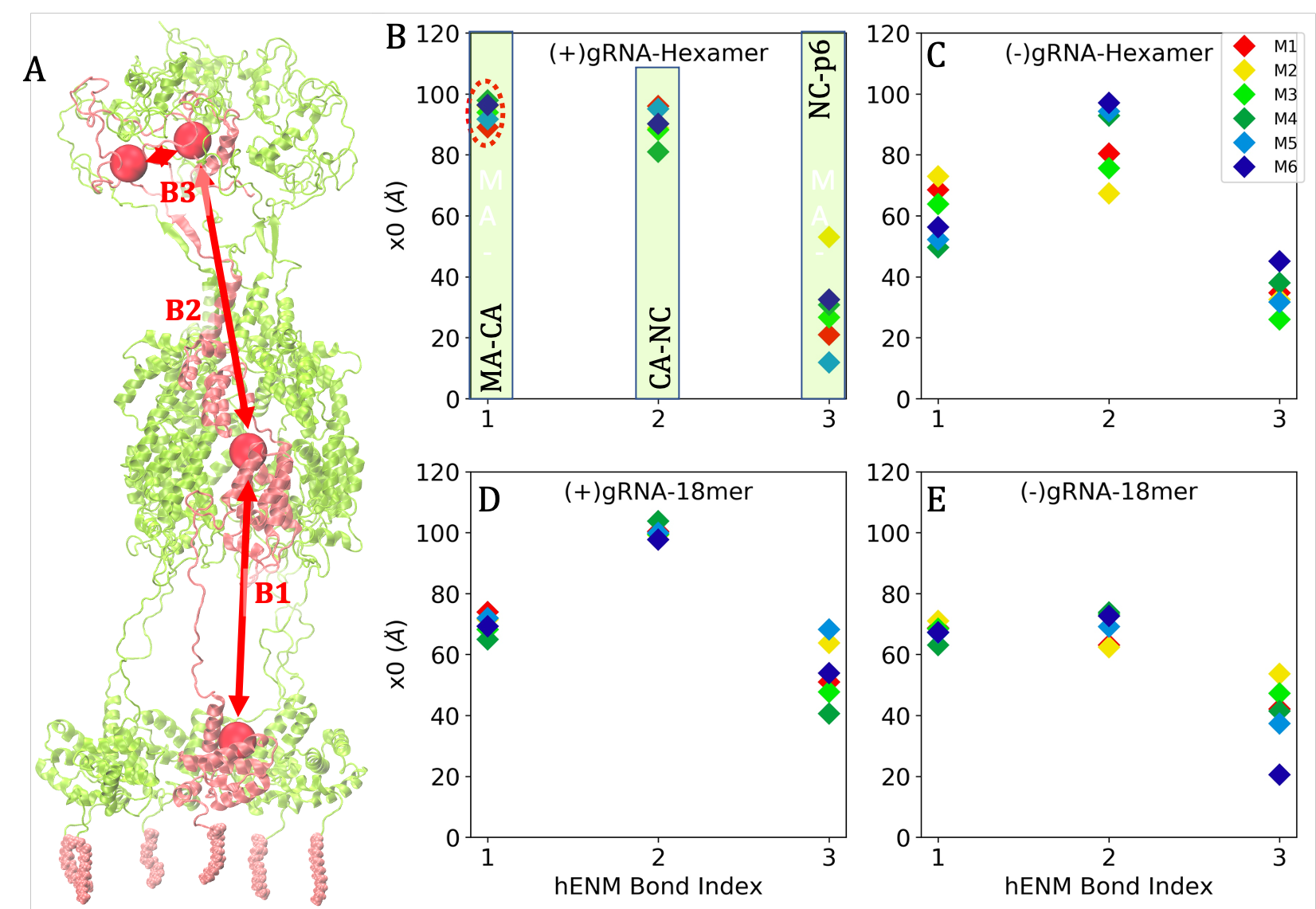

**Figure S2: Equilibrium separation of domains (x_0_ in Å).** A. The construction of the four-site CG model for the hexameric Gag model. (B-C) Equilibrium separation of CG beads for six Gag monomers (M1-6) belongs to hexameric Gag models, with and without implicit gRNA binding, respectively. (D-E) x_0_ values for six Gag monomers in the 18mer Gag model constituting the inner hexameric ring.

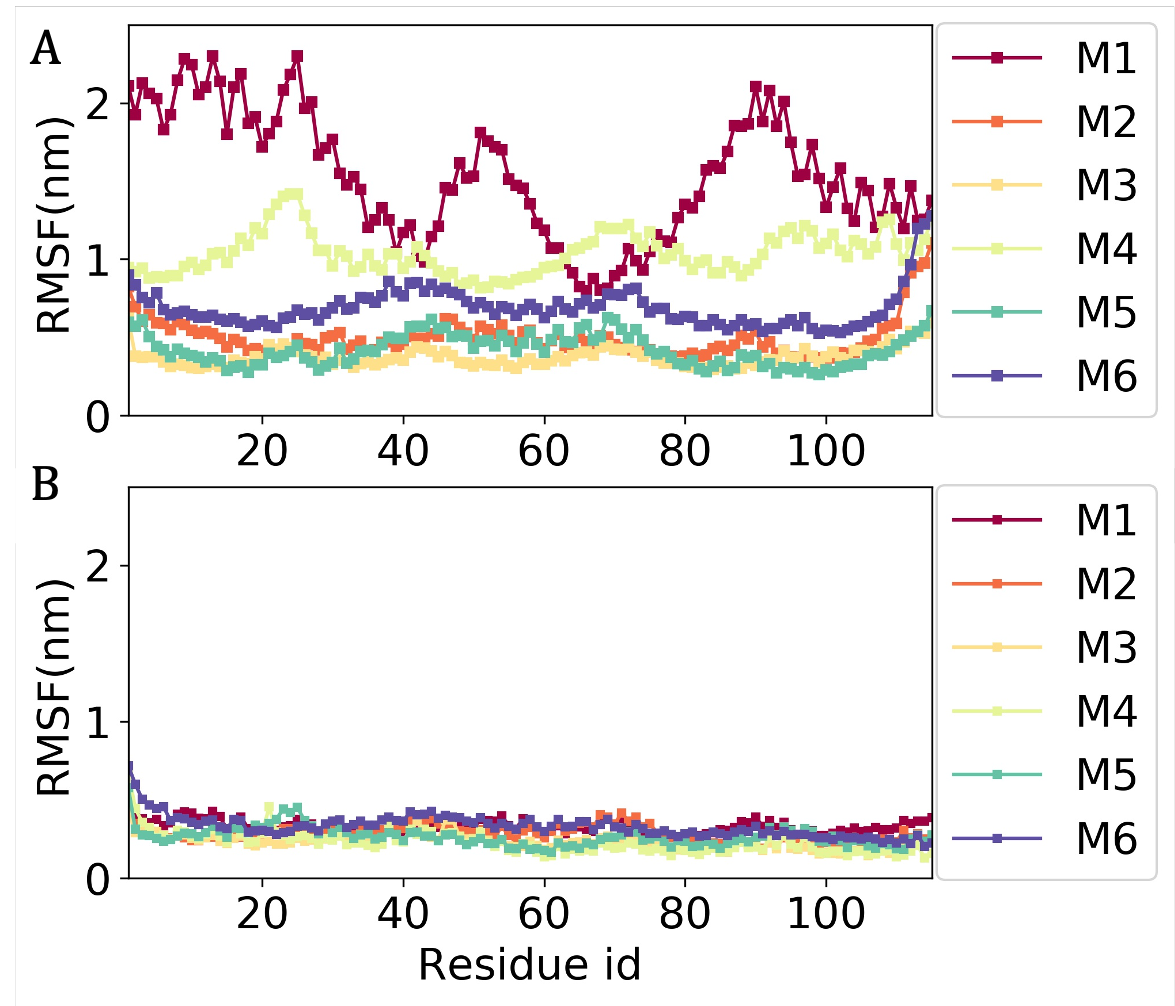

**Figure S3:** **Root mean square fluctuation (RMSF) of MA domain in the membrane-bound hexameric Gag.** RMSF of MA domain (residue 1 to 131) A. with implicit gRNA binding, B. No gRNA binding to NC domain.

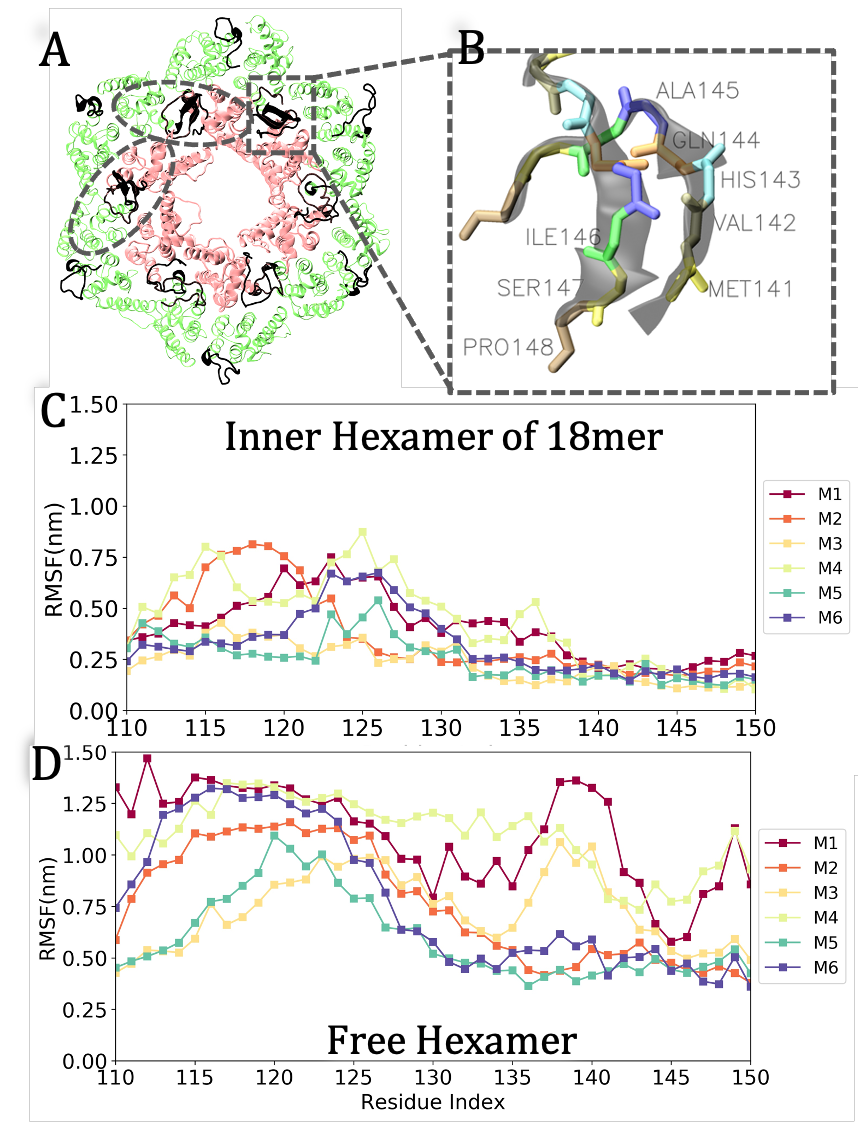

**Figure S4:** **Dimeric interface at CA_NTD_ domain:** A. CA_NTD_ domain of immature Gag 18mer. The inner hexamer is shown in pink and the outer 12 monomers are shown in green. N-terminal residues of the CA domain, residues 131-148 participating in the dimeric interactions are highlighted in black. B. Residues MET141-PRO148 involved in stable anti-parallel β-sheet interactions between two CA_NTD_ neighbors. C-D. Root mean square fluctuation (RMSF) of the MA-CA linker domain of six Gag monomers (forming the inner hexameric ring) of the Gag 18mer and free hexamer model, respectively.

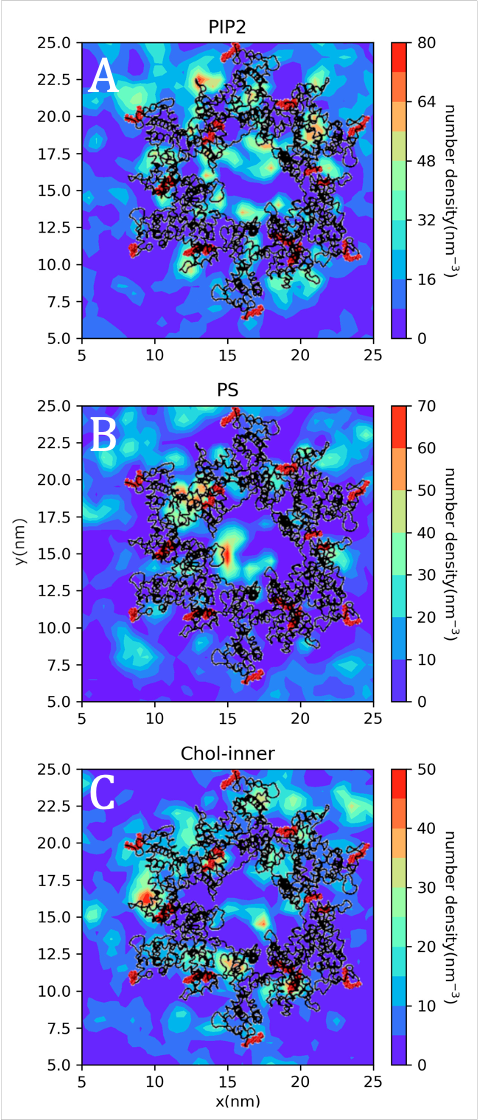

**Figure S5:** **Lipid density map around MA 18mer ((+) gRNA).** The Number density of lipid headgroup atoms at the inner leaflet of bilayer A. PIP2 lipid, B. PS lipid, C. Cholesterol in the inner leaflet. MA protein is shown in black with Myr groups in red.

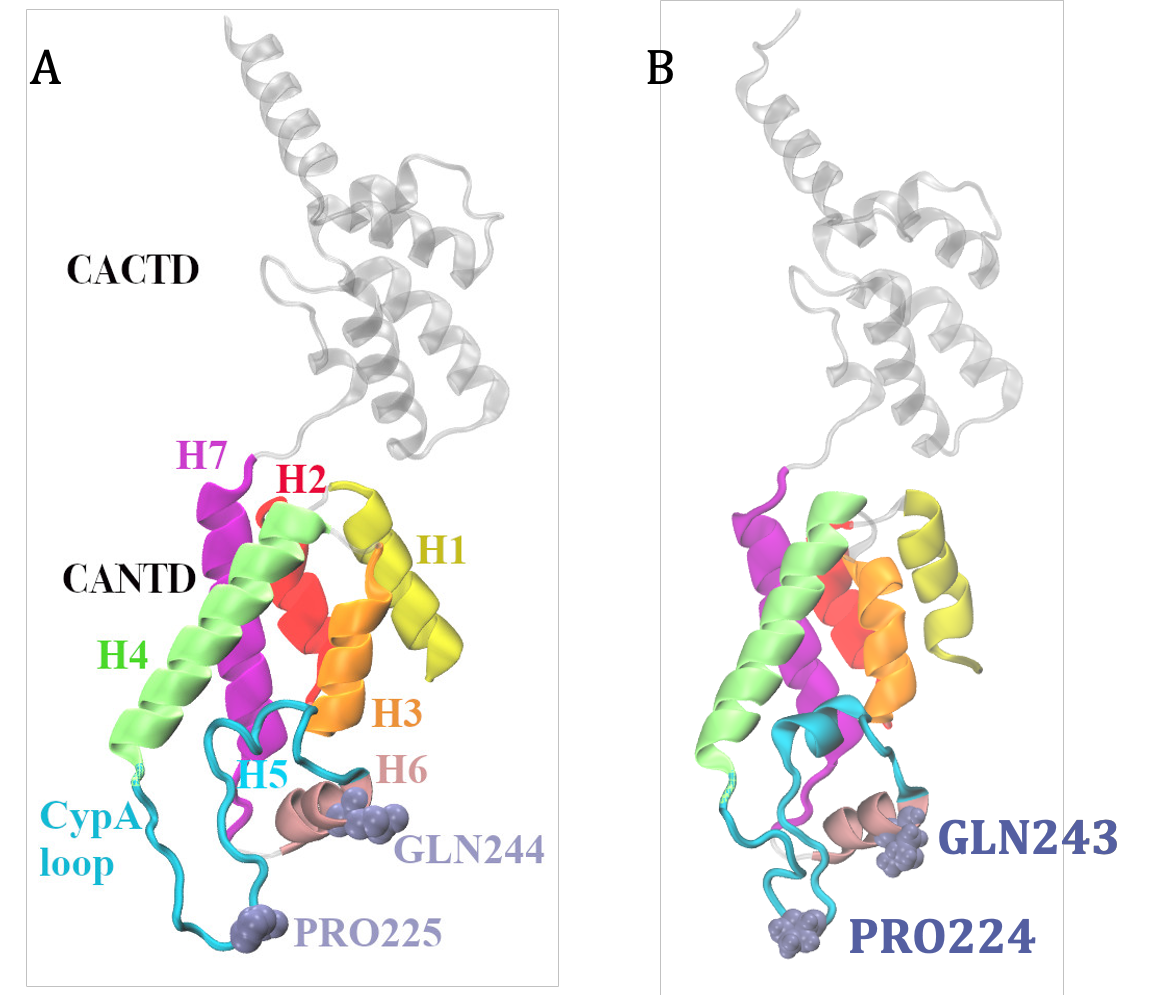

**Figure S6:** **Structure of CA_NTD_ monomer inside Gag lattice structure.** A. CryoET structure [PDB ID: 5L93]. B. Simulated structure of a CA monomer belongs to the inner hexameric ring of the 18mer Gag. Key regions of the CA_NTD_ domain are labeled: Helices 1-6 (in order of appearance in the protein sequence) and CypA loop connecting H4 and H5. GLN243 of H5 and PRO224 of CypA loop (highlighted in **Figure 3**) are shown. Residue_n_ of this Cryo-ET structure is residue_n+1_ of our modeled structure, as in our simulated model residue1 is composed of myristoyl group and GLY1 residue.

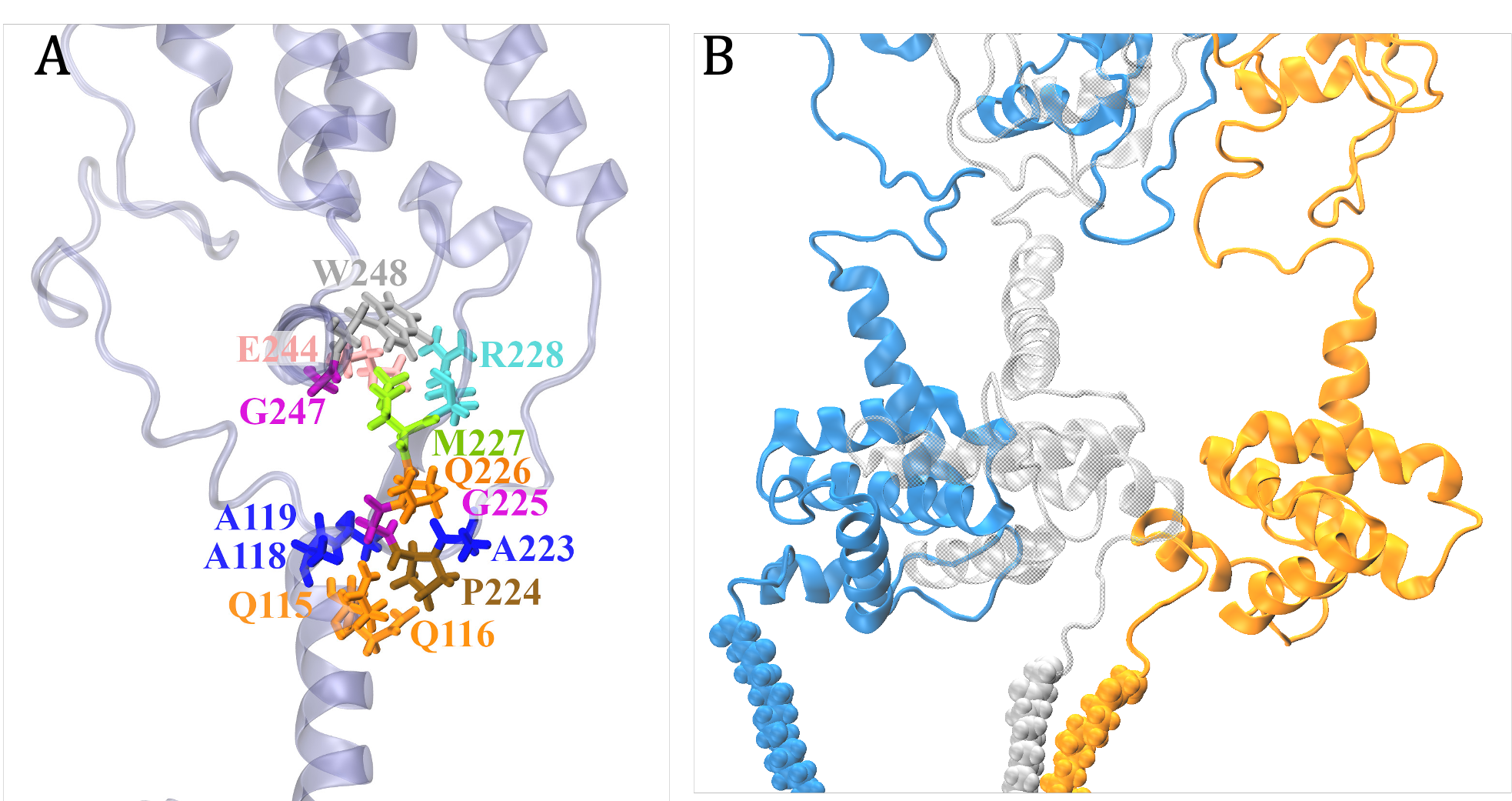

**Figure S7:** **Intra and Interchain interactions mediated through CypA binding loop domain (residue 218-226).** A. CypA domain interacting with MA domain residues in a Gag monomer, part of the inner hexameric ring of Gag 18mer. B. CypA loop of the Gag monomer (shown in blue) interacts with MA domain residues (shown in grey) and transiently interacts with the MA-CA linker domain of the orange Gag monomer. Gag monomers shown in blue and orange comprise the inner hexameric ring of Gag 18mer, whereas the one shown in grey is from the peripheral Gag monomers, which share a dimeric interface with the orange Gag monomer.

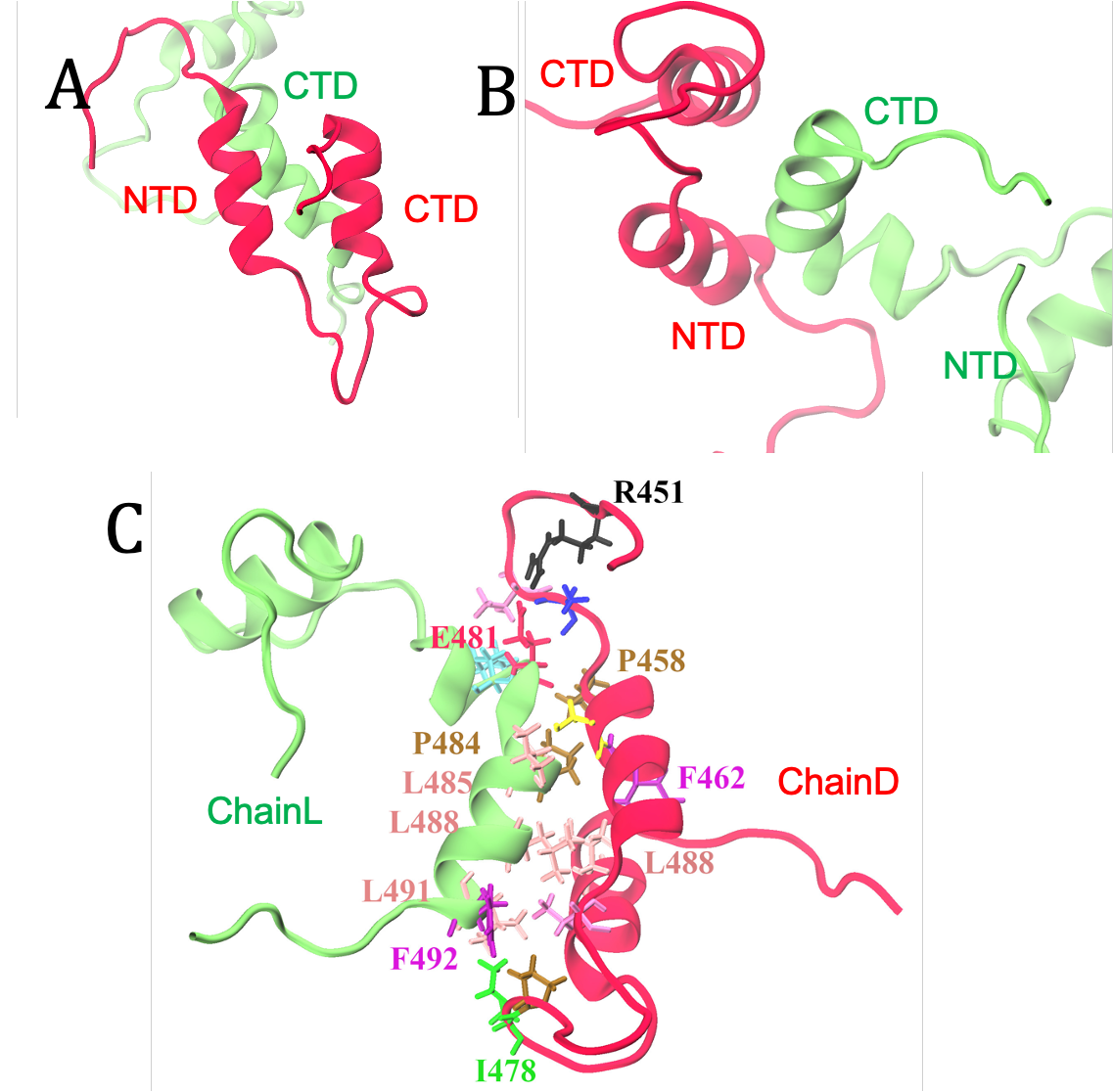

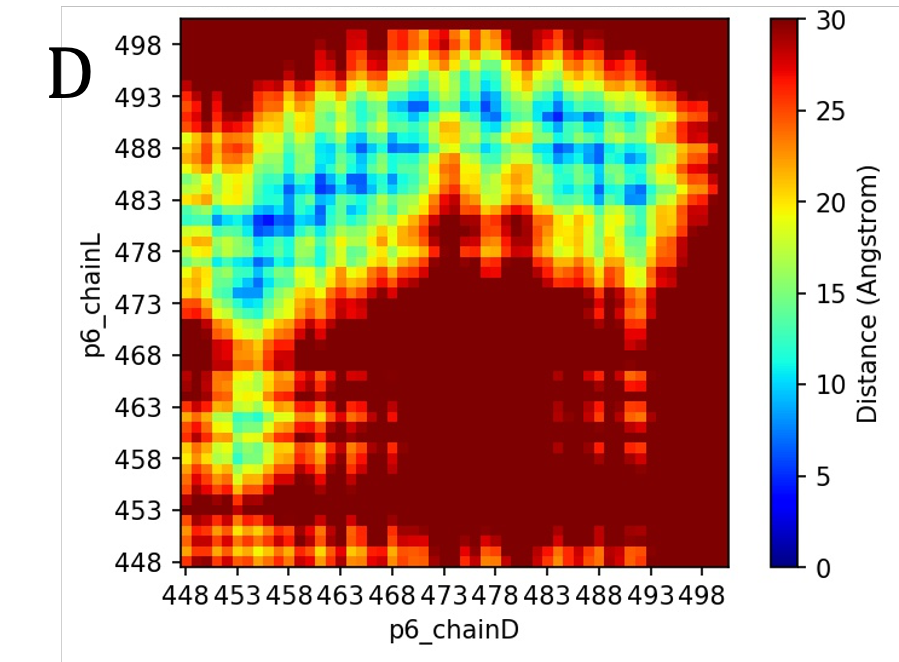

**Figure S8:** **Homo domain interactions mediated by p6. (**A-B) Binding of CTD helix of p6 domain (Chain L, green) to both the helices of ChainD p6 domain. Chain D is a part of the inner hexameric ring of Gag 18mer and Chain L is one of the 12 outer Gag monomers. They do not form any homo-domain interactions except for the p6 domain. C. Such p6-p6 interaction is mediated mainly by hydrophobic amino acid residues. D. Distance map between p6 residues of two chains.
